# Poisson balanced spiking networks

**DOI:** 10.1101/836601

**Authors:** Camille E. Rullán Buxó, Jonathan W. Pillow

**Affiliations:** Center for Neural Science, New York University; Princeton Neuroscience Institute & Dept. of Psychology, Princeton University

## Abstract

An important problem in computational neuroscience is to understand how networks of spiking neurons can carry out various computations underlying behavior. Balanced spiking networks (BSNs) provide a powerful framework for implementing arbitrary linear dynamical systems in networks of integrate-and-fire neurons (Boerlin et al. [1]). However, the classic BSN model requires near-instantaneous transmission of spikes between neurons, which is biologically implausible. Introducing realistic synaptic delays leads to an pathological regime known as “ping-ponging”, in which different populations spike maximally in alternating time bins, causing network output to overshoot the target solution. Here we document this phenomenon and provide a novel solution: we show that a network can have realistic synaptic delays while maintaining accuracy and stability if neurons are endowed with conditionally Poisson firing. Formally, we propose two alternate formulations of Poisson balanced spiking networks: (1) a “local” framework, which replaces the hard integrate-and-fire spiking rule within each neuron by a “soft” threshold function, such that firing probability grows as a smooth nonlinear function of membrane potential; and (2) a “population” framework, which reformulates the BSN objective function in terms of expected spike counts over the entire population. We show that both approaches offer improved robustness, allowing for accurate implementation of network dynamics with realistic synaptic delays between neurons. Moreover, both models produce positive correlations between similarly tuned neurons, a feature of real neural populations that is not found in the original BSN. This work unifies balanced spiking networks with Poisson generalized linear models and suggests several promising avenues for future research.

## 1 Introduction

The brain carries out a wide variety of computations that can be implemented by dynamical systems, from sensory integration [2–5], to working memory [6–8], to movement planning and execution [9–11]. Although the existence of such computations in the brain is well established, the mechanisms by which these computations are implemented in networks of neurons remains poorly understood. One approach to this problem involves statistical modeling, which uses descriptive statistical methods to infer the dynamics of neural activity from recorded spike trains [11–22]. A second approach involves theoretical modeling, which seeks to identify strategies for implementing dynamical systems with networks of idealized model neurons [3, 23–31]. An important example of this second approach is the balanced spiking network (BSN) framework introduced by Boerlin et al [1].

The BSN model consists of a network of coupled leaky integrate-and-fire (LIF) neurons that can emulate an arbitrary linear dynamical system (LDS). The motivating idea is to design a network that approximates the output of a target LDS with a weighted combination of filtered spike trains. The population is divided into “excitatory” and “inhibitory” populations of neurons, based on whether they contribute positively or negatively to the output. This leads to an intuitive spiking rule: a neuron should spike whenever doing so will reduce the error between the output of the target LDS and the network output, i.e., the weighted combination of filtered spikes emitted so far. To make this work, each neuron has to maintain an internal representation of the error between the desired LDS output and the current network output. Boerlin et al showed that, remarkably, this computation can be mapped precisely onto the dynamics of an LIF neuron. A neuron’s membrane potential is a local representation of the network-wide error between target output and current network output, and its spike threshold is proportional to the amount by which adding a spike will reduce this error.

The BSN framework has many appealing characteristics. Spiking is efficient, in the sense that every spike contributes meaningfully to reducing error between target and actual output. The computations performed by the model are robust to perturbations and to the loss of neurons. The model also generates irregular spiking activity with intervals that that resemble those observed in real neurons.

However, the original BSN model has an important shortcoming that limits its plausibility as a model for information processing in real neural circuits. Namely, the model requires unrealistically fast propagation of information between neurons. Because every neuron’s membrane potential is tracking the overall error between target and actual output, the membrane potential of all neurons has to reset whenever any neuron emits a spike. Failure to impose this reset leads to increased activity as multiple neurons attempt to correct same error. In fact, implementations of the BSN model typically impose a rule enforcing that only one neuron is allowed to spike in a single time bin, effectively allowing spikes to propagate faster than the temporal resolution of the simulation (e.g., 0.1ms). Without this rule, the network can easily enter unstable modes in which excitatory and inhibitory populations emit massive spike bursts in alternating time bins, overshooting the target in an attempt to correct the error from the previous time bin.

Here we show that a probabilistic spiking rule can overcome the need for unrealistically fast propagation of spikes in the BSN framework. The basic intuition for our solution is that instead of making neurons spike deterministically whenever doing so will reduce error, we can allow multiple neurons to spike probabilistically such that error will be reduced on average.

We propose two alternate formulations of BSN with Poisson spiking, distinguished from each other by the level at which the network is attempting to minimize the decoding error. First, we describe a ‘local’ framework, which preserves the original BSN model dynamics but replaces the hard integrate-and-fire spiking rule with the soft firing threshold of the Poisson generalized linear model (GLM) [32–35]. This spiking rule generates stochastic spiking conditioned only on each neuron’s local copy of the error which, on average, leads to a reduction in the population-level read-out error.

Second, we propose a ‘population’ framework that replaces the greedy, single-neuron perspective of the local and BSN models with a rule based on minimizing the expected error at the population level. A vector of spike rates is generated by calculating the expected spike counts that minimizes the total decoding error. The probability of a single neuron spiking depends on its own weight, as with the local rule, but also takes into account the activity of the entire population of neurons and their weights. This coordination leads to spiking activity that is efficient and invariant to network size. Finally, we show that both the local- and population-level Poisson frameworks make the BSN robust to synaptic delays.

This paper is organized as follows. We begin with a pedagogical review to the BSN model (Sec. 2). We then examine the model’s dependence on instantaneous spike propagation, and document the unstable behavior that arises if multiple spikes are allowed in a single time bin (Sec. 3). To address this shortcoming, we introduce local and population BSN models with conditionally Poisson spiking (Sec. 4). Finally, we illustrate the accuracy and robustness of these models to synaptic transmission delays (Sec. 5).

## 2 Background: balanced spiking network model

Here we provide a brief introduction to the original balanced spiking network (BSN) framework introduced by Boerlin, Machens, & Denève [1]. The goal is to design a spiking network that can accurately implement an arbitrary linear dynamical system. Consider a linear dynamical system defined by:

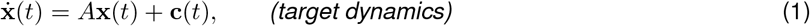

where **x**(*t*) = (*x*_1_(*t*)…*x_J_*(*t*))^⊤^ is a vector of *J* dynamic variables that we will refer to as the *target, A* is the *J × J* linear dynamics matrix, and **c**_*t*_ = (*c_i_*(*t*), …*c_J_*(*t*))^⊤^ is a *J*-dimensional vector of inputs. The BSN model consists of a spiking network of *N* neurons that attempts to approximate the target output **x**(*t*) via a weighted combination of filtered spike trains:

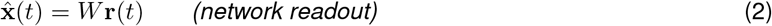

where **r**(*t*) is the set of spike trains convolved with an exponential decay function, and *W* are *J* × *N* readout weights. (See Fig. 1 for a schematic.) In general,fora 1-D dynamical system,the population is divided into equal pools of ‘positive’ and ‘negative’ neurons (depending on the signs of their individual weight components)although this is not a strict requirement. For *J* > 1, the ‘positive’ vs ‘negative’ distinction does not necessarily apply as the signs of the weights need not be consistent across dimensions.

**Figure 1:**
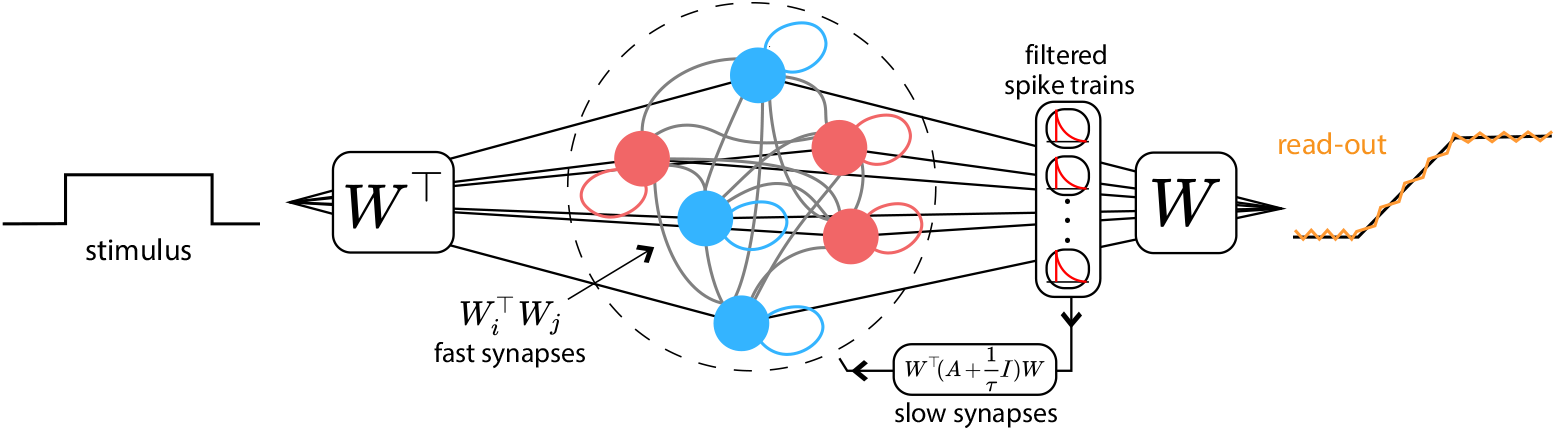
A simple diagram illustrating the BSN. Neurons receive stimulus input projected onto the transpose of a set of linear weights, *W*^⊤^, and the output is reconstructed by filtering spikes through the same weights, *W*. Neurons are connected via two coupling weights: fast synapses, *W*^⊤^*W*, which instantaneously propagate individual spikes through the network, and slow synapses, 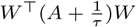, which implement network dynamics by feeding the filtered spike trains back into all neurons in the network. The network is divided into two equal populations of positive (red) and negative (blue) output weights, whose spikes have opposite effects on network output.

The *i*’th component of the vector **r**(*t*) is given by

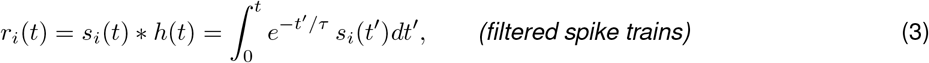

where 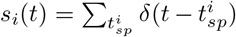 denotes the *i*’th neuron’s spike train, defined by a series of delta functions at spike times 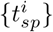, and *τ* is the time constant of the exponential filter *h*(*t*).

From this starting assumption, Boerlin *et al* introduce a greedy update rule that causes a neuron to spike whenever doing so will reduce the squared error between target **x**(*t*) and network output 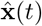,

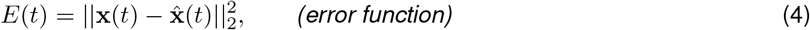

which is mathematically equivalent to the threshold-crossing spiking rule in a leaky integrate-and-fire (LIF)neuron.

Here we recapitulate the derivation of this spiking rule in discrete time, for clarity and ease of implementation. Let 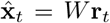 denote the network output at time bin *t*. The effect of adding a spike from neuron *i* in this time bin would be to augment the output vector by that neuron’s decoding weight vector **w**_*i*_, which is given by *i*’th row of the decoding weight matrix *W*. Thus, network output is 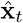 if neuron *i* is silent and 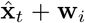 if it spikes. This suggests that the neuron should spike if doing so will result in smaller error, or

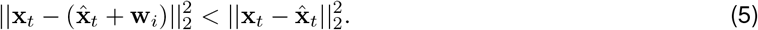

Simplifying this expression yields the condition that the neuron should spike if projection of the error vector onto **w**_*i*_ is greater than half the squared *L*_2_ norm of **w**_*i*_:

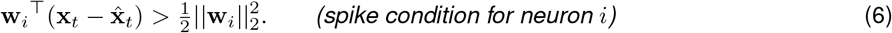

Boerlin *et al* therefore suggest regarding the time-dependent left hand side of (eq. 6) as the membrane potential for neuron *i*, and the right hand side as its spike threshold *T_i_*:

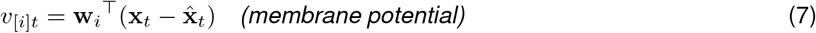

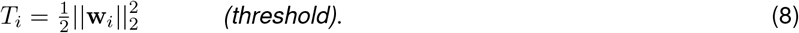

Under this view, each neuron is computing a local approximation to the difference between the true target **x**_*t*_ and the network’s current output 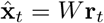, projected onto that neuron’s weight vector **w**_*i*_.

The only missing piece from this expression is that the neurons do not of course have access to the true value of **x**_*t*_. But they do have implicit access to *A*, and thus to the dynamics governing **x**_*t*_. Boerlin et al therefore propose that the the network output 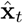 is sufficiently close to the target output **x**_*t*_ that it can be used to accurately approximate the desired dynamics: 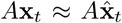. To make this explicit, we introduce a proxy variable **z**_*t*_, which denotes the network’s own (internal) approximation to the true target **x**_*t*_. This proxy variable evolves according to

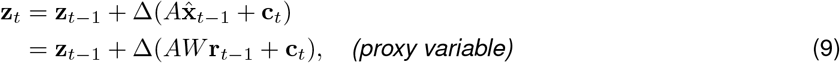

where for simplicity we use a forward Euler method for integrating the dynamics equation (eq. 1) with time bin size Δ. Higher accuracy can be achieved using exponential Euler integration (see Methods). Of course **z**_*t*_ is never represented explicitly; the network tracks **z**_*t*_ via its projection onto the decoding weights W, as we will see shortly.

### 2.1 Simulating the BSN model

Simulating the BSN model for a single time bin can be described by a sequence of three steps:

1. Calculate the “pre-spike” membrane potential for each neuron by combining inputs from the previous time step and external input.
2. Apply the threshold to determine which neurons (if any) emit spikes.
3. Reset to obtain “post-spike” membrane potentials **v**_*t*_ and update filtered spike trains **r**_*t*_.

We will describe each of these steps in turn. First, the update rule for the pre-spike membrane potential (consistent with eq. 7) is:

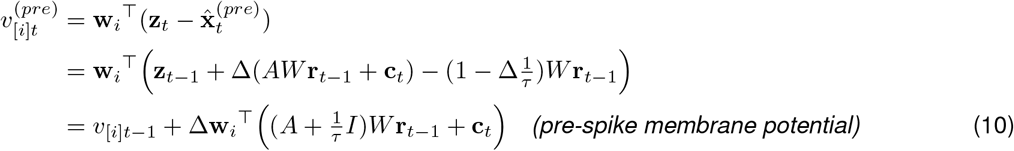

where 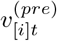 denotes the pre-spike membrane potential for neuron *i* at time bin *t*, 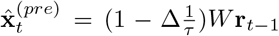 denotes the network output for the current time bin *before spiking*, and *v*_[*i*]*t*−1_ = **w**_*i*_^⊤^(**z**_*t*−1_ − *W***r**_*t*−1_) denotes the (post-spike) membrane potential from the previous time step.

Second, spikes for the current time bin are computed by determining whether pre-spike membrane potential exceeds threshold 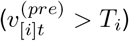. When this occurs, the neuron records a spike: *s*_[*i*]*t*_ = 1.

Lastly, the filtered spike trains *r*_[*i*]*t*_ are augmented and membrane potential is reset:

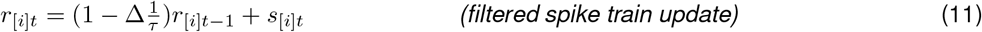

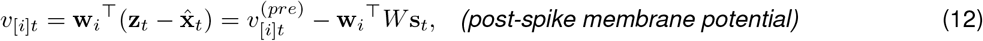

which ensures that post-spike membrane potential equals the difference between the projected proxy variable and network output.

### 2.2 Vector update rules

For convenience, we can rewrite the BSN update equations in vector form. The pre-spike membrane potential is given by:

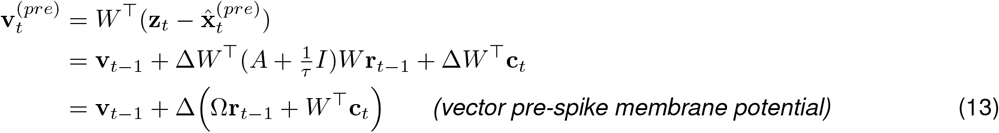

where 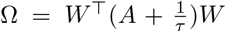 are the coupling weights from **r**_*t*−1_ to the pre-spike membrane potential, which implement computation of the divergence between the target **z**_*t*_ and passive decay of **r**_*t*_ in the absence of spiking.

Once the spike vector **s**_*t*_ has been computed, the filtered spike trains and network output for the current time bin are given by:

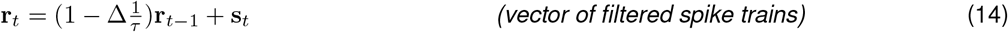

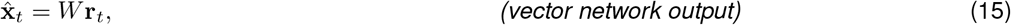

and the vector of post-spike membrane potentials is given by

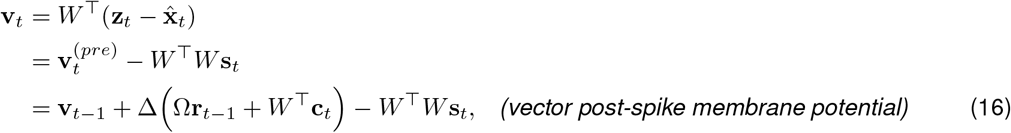

which reflects reset of the pre-spike membrane potential after spiking, but can equivalently be seen to be the projected difference between the proxy variable **z**_*t*_ and the current network output 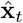 in the current time bin.

It is worth noting that this model requires the instantaneous propagation of spikes between neurons. After a spike, the membrane potential reset (eq. 16) updates **v**_*t*_ for all neurons based on the spikes in the current time bin via the fast weights, −*W*^⊤^*W*. Although Boerlin *et al* refer to the weights −*W*^⊤^*W* as “fast synapses” and the Ω as “slow synapses”, note that the Ω**r**_*t*−1_ term also involves near-instantaneous propagation of information, since the exponentially-filtered spike trains **r** jump by 1 after every spike.

The full BSN model first described in [1] contained additional penalties on **r**_*t*_ in the objective function, which had the effect of reducing spiking by trading off minimization of error (eq. 4) against a cost of inserting spikes. Although we have left these terms out our derivation here for simplicity, including them has the limited effect of changing the spike threshold and post-spike reset and does not change the nature of our findings. The simulations shown in the following sections use the full version as described in the Methods section (8). Simulation parameters for each figure are also included in section (8).

## 3 Limitations of the BSN model

A key limitation of the BSN model is that it requires unrealistically fast communication between neurons. In the standard integrate-and-fire model, a spike resets only the membrane potential of the neuron that emitted it. In the BSN model, by contrast, the membrane potentials of *all* neurons reset following a spike from any neuron via the −*W*^⊤^*W***s**_*t*_ term in (eq. 16). The instantaneous reset of all membrane potentials following a spike is necessary to ensure that each neuron’s membrane potential maintains an accurate representation of the read-out after each spike. From a normative standpoint, the hard LIF threshold entails that maintaining an accurate local copy of the error is critical for the network to encode the target.

In fact, the problem is slightly more complicated: standard implementations of the BSN model include a constraint that only one neuron is allowed to spike in a single time bin. Without this constraint, neurons with similar output weights tend to spike in the same time bin, when a spike from any one of them would have sufficed to compensate for error in network output. This causes the network output to dramatically overshoot the target. In the subsequent time bin, neurons with opposite-sign readout weights fire to compensate for this error, and overshoot the target by a large amount in the opposite direction. This sets up a pathological pattern of oscillatory firing known as “ping-ponging”, in which two populations spike maximally in alternating time bins of the discrete simulation [1].

Fig 2 illustrates how ping-pong behavior can arise if multiple spikes are allowed in a single time bin. We set the BSN model to implement a 1-dimensional perfect integrator, *ẋ*(*t*) = *c*(*t*), using the same parameters as the example from figure 1C of Boerlin et al. [1]. In brief, the network contained 400 neurons, divided into two equal sized populations with output weights of +0.1 and −0.1, respectively, which we refer to as positive-output and negative-output neurons (see Methods for complete details). When the rule forbidding multiple spikes per time bin is imposed, the network accurately tracks the target output variable (Fig. 2A). However, removing this constraint — allowing all neurons with membrane potential above threshold to fire — results in ping-pong behavior and large errors in tracking the target (Fig. 2B).

**Figure 2:**
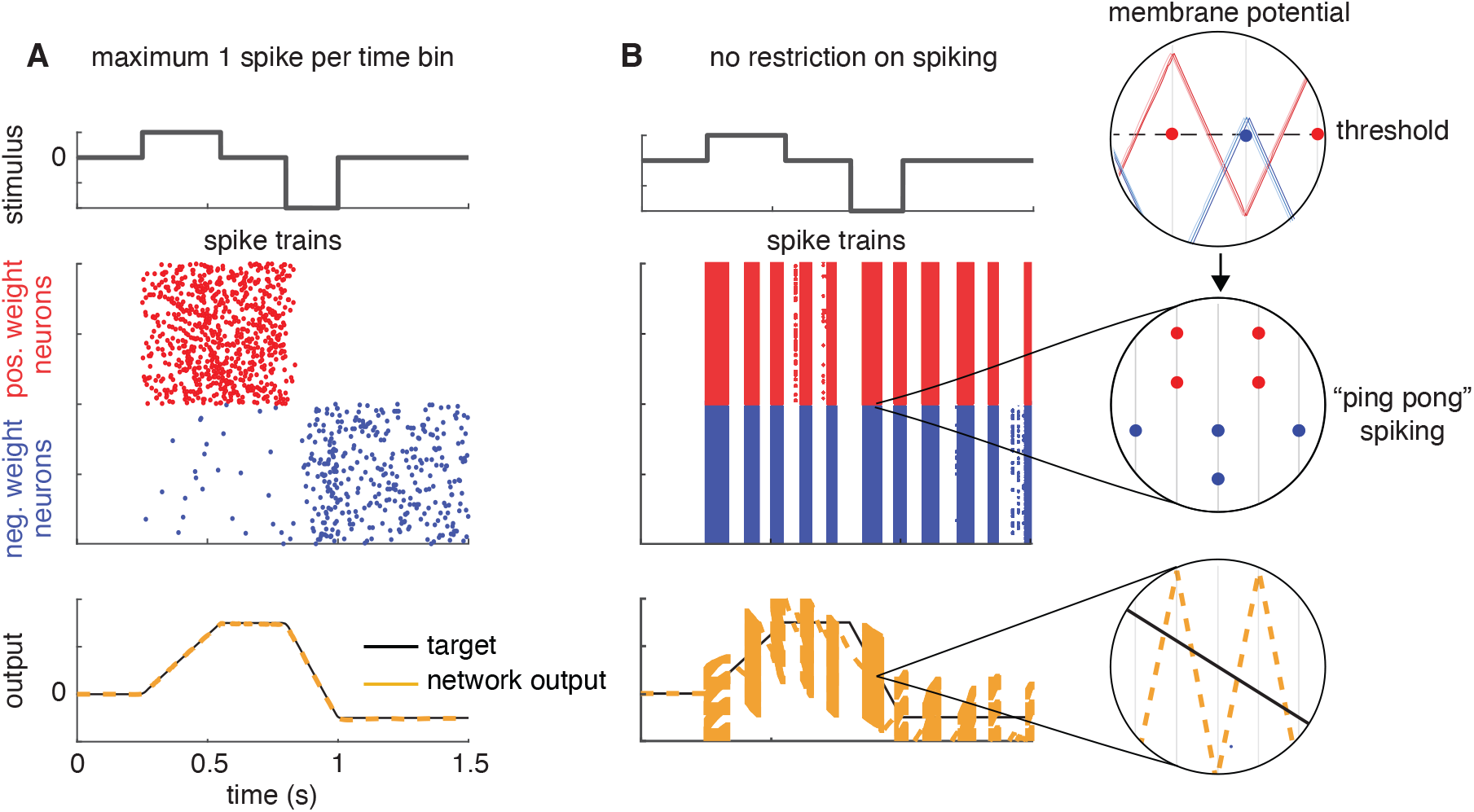
Balanced spiking network implementing a perfect integrator. The network consists of 400 neurons, divided into two populations with output weights of +0.1 (red) and −0.1 (blue). **(A)** Simulation results under the condition that only one neuron is allowed to fire per discrete time bin. **(B)** Simulation results when all neurons whose membrane potential is above threshold in a single time bin are allowed to fire, leading to “ping-pong” behavior. Insets show that read-out (yellow) is alternating between large over- and under-estimates of the target (in black). Inset shows the voltage traces of neurons in both positively and negatively weighted populations. Since the weights and inputs are identical across populations, so are the voltage traces. As a consequence, during ping-ponging, all neurons within a population cross the threshold in the same time bin.

Ping-ponging can in principle be eliminated by adding noise to the membrane potential and using extremely small time bins, which increases the probability that at most one neuron will cross threshold in a single time bin. However, we found that this required a reduction in time bin size by several orders of magnitude. Moreover, this solution relies even more heavily on instantaneous synaptic communication, since it allows neurons to reset even more rapidly after a spike from any single neuron.

## 4 BSN with conditionally Poisson neurons

To overcome the problems of instantaneous transmission and network instability, we propose two novel formulations of balanced spiking networks with conditionally Poisson neurons: (1) a local framework, where each neuron spikes independently based on its local estimate of network error; and (2) a population-level framework, which sets firing rates to reduce expected error for the entire population.

The key idea in both frameworks is to replace the deterministic integrate-and-fire spiking rule with probabilistic spiking. Under this modified spiking rule, spiking is governed by an instantaneous probability of spiking λ_*t*_, also known as the *conditional intensity*, such that spiking is independent with probability Δλ_*t*_ in any small time window of width Δ. This results in an auto-regressive Poisson generalized linear model (GLM), also known as a Cox process [33–35]. This model has a quasi-realistic biophysical interpretation [32, 36, 37], and recent work has shown that it can capture a wide range of dynamical behaviors found in real neurons [35].

### 4.1 Local framework

A simple way to introduce probabilistic spiking to the BSN framework is to replace the hard spike threshold of the integrate-and-fire model with a soft threshold, so that spike probability grows as a nonlinear function of membrane potential, an approach also known as the “escape-rate approximation”[32]. Specifically, we define each neuron’s conditional intensity function to be a sigmoidal function of membrane potential:

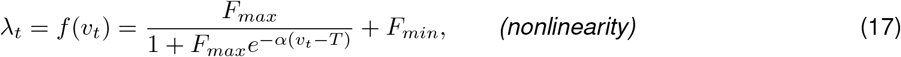

where *v_t_* is the membrane potential at time *t, T* is the spike threshold, *F_max_* is the maximal firing rate, *F_min_* is a baseline firing rate, and *α* is a slope parameter governing the sharpness of the threshold. The probability of spiking in a small time interval is proportional to λ_*t*_, which models the spike response as an inhomogenous Poisson process. We refer to this as the local Poisson framework because, like the BSN, spikes are generated by internal dynamics that evolve according to local copies of the representation error.

Although it is common to use exponential nonlinearities for Poisson GLMs, here we have used a scaled sigmoid function to control both the suddenness of firing onset and the maximum achievable firing rate. The parameter *α* controls the precision of firing onset, while *F_max_* and *F_min_* control the range of firing rates within a small time window. The resulting function (eq. 17) resembles an exponential function at low firing rates but saturates at a maximum of *F_max_* (see Fig. 4). If we let both *α*, *F_max_* → ∞, we recover the original (hard-threshold) integrate-and-fire rule, in which a spike occurs probability 1 when *v_t_* > *T*. However, for finite *α* and *F*, the onset of spiking is more gradual.

As a practical matter, we do not wish to allow a single neuron to fire multiple spikes in a single time bin, because the first spike would preclude additional spikes in the same time bin due to “reset” of the membrane potential. We therefore simulate the model with the spiking rule:

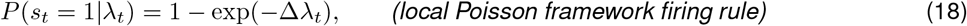

where exp(−Δλ_*t*_) is the probability of observing no spikes in a time bin of size Δ under the Poisson model. For each time-step, we update **v** as in the original BSN model, pass it through the nonlinear function *f*(·) to obtain the vector of Poisson firing rates, λ_*t*_, and draw spikes as independent Bernoulli random variables with probability as given above. See Fig. 3 for a schematic.

**Figure 3:**
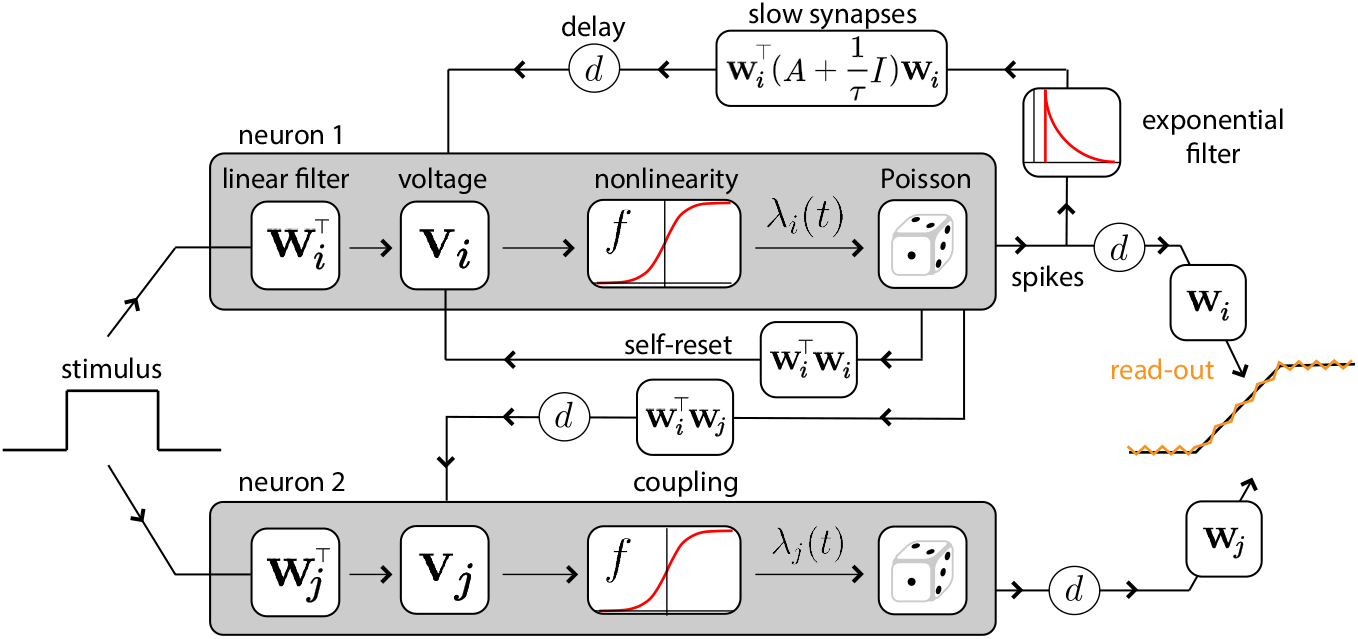
Schematic of BSN with conditionally Poisson neurons. The stimulus influences each neuron’s membrane potential **v**_*i*_ via a set of input weights *W*^⊤^. The neurons reset themselves via instantaneous, fast synapses. Fast connections to other neurons propagate the effects of spikes with a synaptic time delay *d*. The desired linear dynamics are implemented via slow weights (filtered through an exponential) also with a time delay *d*. Within each neuron, spiking is probabilistic with an instantaneous probability of firing λ_*i*_(*t*) = *f*(*v_i_*(*t*)), where *f*(·) is a nonlinear function of voltage.

Fig. 4D shows an illustration of how *α* affects spiking precision and Fig. 4E shows a comparison between spike trains generated by a single-neuron integrating a noisy stimulus for different values of *α* and those of the deterministic LIF model (in black). As we expect, for high values of *α* we recover the precise spiking behavior of the BSN model. As *α* decreases, spike times spread around the ideal BSN spikes. The sub-threshold voltage traces also change with respect to *α*, indicating that the spiking behavior is a reflection of an underlying error-driven probabilistic process.

**Figure 4:**
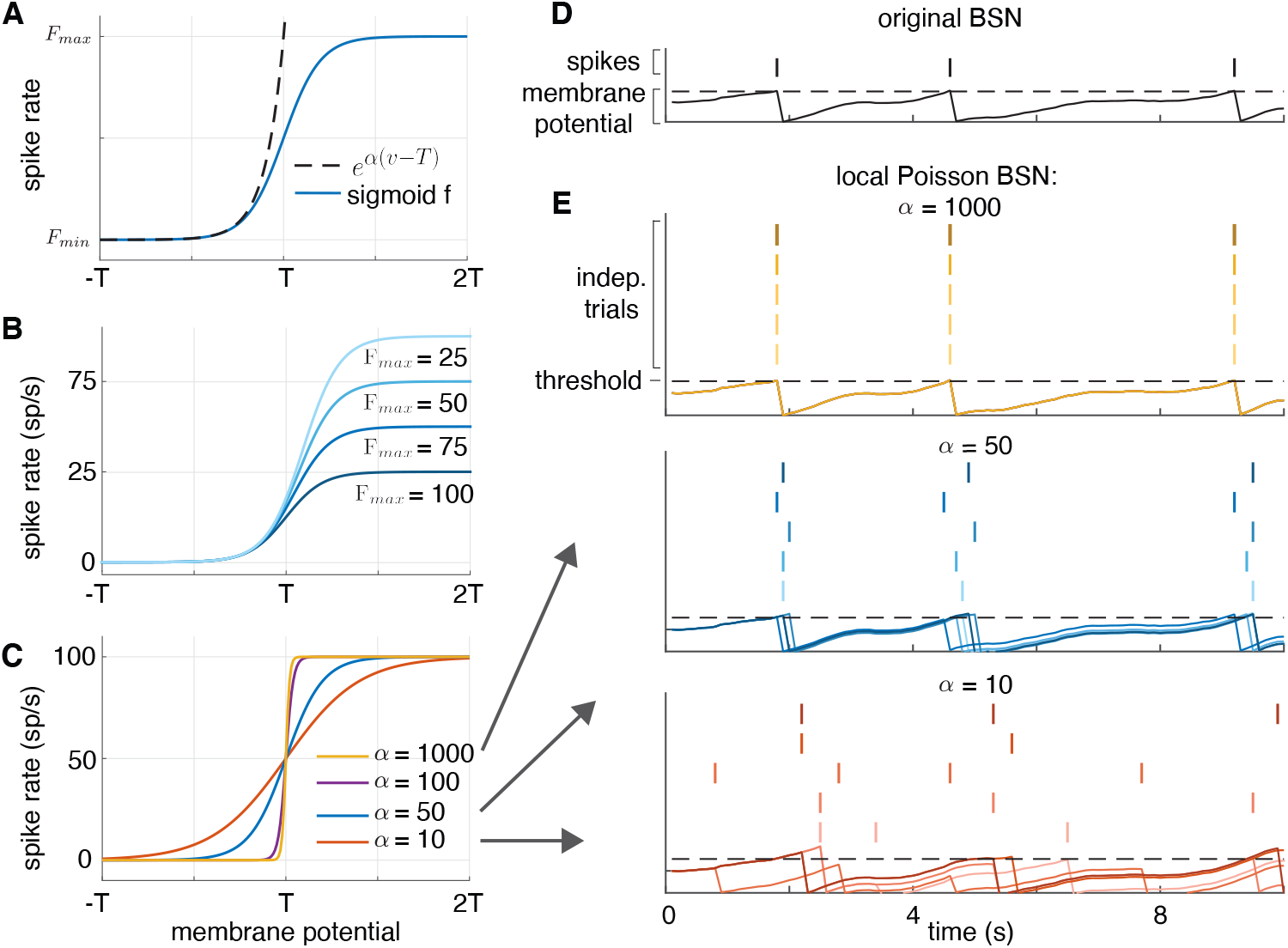
**(A)** The conditional intensity for the exponential non-linearity (dashed lines) and the sigmoid non-linearity (solid lines). The conditional intensity of the sigmoidal non-linearity closely follows that of the exponential non-linearity for sub-threshold voltages, but levels off after threshold, keeping firing rates stable. **(B)** Family of nonlinearities with varying *F_max_*. Increasing *F_max_* raises the firing rate at which the nonlinearity saturates. **(C)** Family of nonlinearities with varying *α*. Increasing *α* increases the steepness of the nonlinearity, which approaches a hard-threshold function as *α* → ∞ (like the BSN). **(D)** Simulation of the original BSN implementing a perfect integrator, showing membrane potential and spikes of a single example neuron. **(E)** Spikes and membrane potential of the same neuron in a local Poisson BSN implementation of the same system. High *α* simulations (yellow) replicate the behavior of the BSN integrator. Lowering *α* to 50 (blue) or 10 (red) results in a spread of spikes centered around the deterministic BSN spikes.

#### Robustness to parameters of the nonlinearity

We studied the effects of varying *α*, *F_max_*, and *F_min_* on the performance of the homogeneous integrator network (5) and found that there exists a wide range of values for which the network error is low and the spiking activity is efficient (5E-F). Figures 5A shows raster plots and corresponding read-outs for ‘optimal’ parameter settings (defined as being in the low error and activity range, denoted by the * in 5E-F), and in 5B-D we modify these parameters to show qualitative changes in spiking activity and read-out accuracy. Decreasing *α* and *F_max_* negatively impacts read-out quality, while removing background spiking returns the network to a high-precision, synchronized regime.

**Figure 5:**
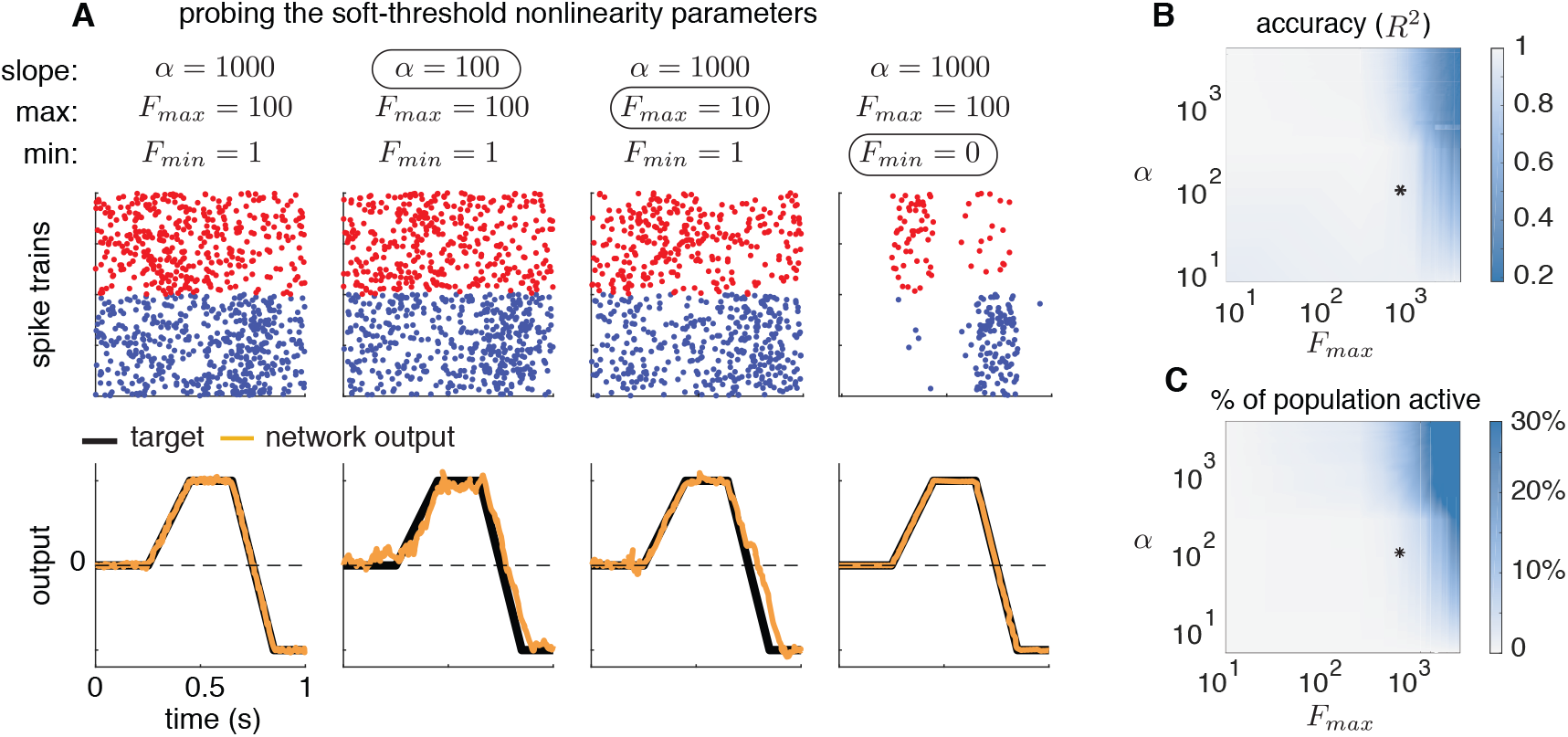
**(A)** Simulations of local Poisson model showing the effects of varying the parameters of the soft-threshold nonlinearity on performance. Relevant parameters are the slope *α*, maximal firing rate *F_max_*, and baseline firing rate *F_min_*. Network dynamics implemented a perfect 1D integrator and the stimulus was the same as Fig. 2. Red and blue dots indicate spikes from neurons with positive and negative output weights, respectively. **(B)** Network performance as quantified by *R*^2^ across a range of parameter settings with baseline fixed at *F_min_* = 1. Black asterisk indicates the values for the rightmost column of A (*α* = 1000, *F_max_* = 100). Accuracy remains high across a broad range of parameter values, falling substantially below 1 when slope and maximum firing rate are both large, which gives rise to ping-ponging. **(C)** Percent of the neural population active as a function of *α* and *F_max_*, showing ping-ponging behavior in upper right corner, where the model approaches a deterministic, hard-threshold firing rule.

Network performance noticeably deteriorates for very large values of *α* and *F_max_* because we are forcing the precision to be too high while allowing neurons to spike too frequently. Since the spiking rule is localized, large proportions of the population are active at the same time (5F), much like the ping-ponging seen in the BSN model. However, the values at which this happens (e.g., *F_max_* > 10^3^ spikes/second) are well above what we would expect to see in a biological system. Although for the sake of clarity we present the simulation for a relatively simple target function, we still observe a wide ranges of stable *F_max_* and *α* settings for more complex or multi-dimensional simulations.

The computational advantage of our probabilistic spiking rule is that it introduces uncertainty into spike timing, therefore preventing all neurons from firing at once. The parameters controlling this asynchrony, *F_max_*, *F_min_* and *α*, have direct physical interpretations (maximal and minimal/background firing rate and error tolerance, respectively) which can be mapped on to characteristics of real neural circuits. By introducing extra degrees of freedom, we are losing the strict normative angle of the original BSN framework. However, what we gain in the process - a considerably expanded space of stable network configurations - makes it possible to introduce realistic communication delays between neurons (see section 5).

### 4.2 Population framework

We now describe a second framework for implementing BSNs with conditionally Poisson neurons. In this approach, we take a population-level instead of a neuron-level view of the optimization problem to be solved. Instead of assigning each neuron to carry an independent representation of the error between the target and actual network output, we assign each neuron an analog probability of firing such that expected number of spikes across the network compensates appropriately for the total error. We refer to this as the population framework.

The derivation of this framework starts from an error function describing the discrepancy between target and actual network output. However, instead of specifying that each neuron should spike whenever doing so will reduce error (eq. 5), we compute a vector of spike rates λ, such that the expected spike response across the population over some time window of length *κ* will minimize error. This leads to the following network objective function:

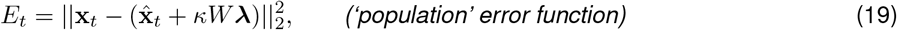

where **x**_*t*_ is the target output at time *t*, 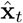 is the actual network output at time *t*, *W* are the decoding weights, and λis the vector of firing rates (conditional intensities) of a network of Poisson spiking neurons. In this expression, *κW*λ is expected contribution to network output over a time window of size *κ*. For implementation in discrete time, *κ* should be an integer multiple of the bin size Δ.

To minimize the above error, we set the instantaneous spike rate vector equal to the least-squares solution:

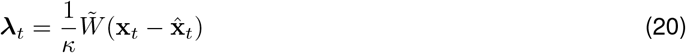

where 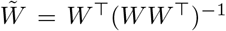 is the Moore-Penrose pseudo-inverse of the decoding weight matrix *W*. Poisson neurons firing independently with conditional intensity λ_*t*_ will therefore minimize the expected error between target and actual network output. Note that if *W* has orthogonal unit-vector rows, such that *WW*^⊤^ = *I*, then 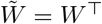 and we obtain the same encoding weights as the original BSN framework.

In the local framework, increasing the population size means more neurons are competing to reduce the read-out error in a single time window. This increased activity can lead to ping-ponging. The population level view of the problem scales the probability of spiking, λ_[*i*]_, by *WW*^⊤^, which increases with population size. The responsibility of correcting an error is thus spread across the entire population and activity is stable with respect to network size.

However, the solution in (eq. 19) is not valid generally because the right-hand-side can take on negative values, whereas the conditional intensity for a Poisson process must be positive. To overcome this, we create two mirrored copies of the population. Positive firing rates are assigned to one copy with weight vector *W*, and negative firing rates are assigned to the other copy with weight vector −*W*. We distinguish between these neurons and ‘anti-neurons’ by the sign of their membrane potential as determined by the least squared solution. Similarly to the local and BSN models, spikes from either population will have opposite effects on the output variable. However, in this case these designations are not fixed labels and do not apply to the actual sign of **w**_*i*_. For example, spikes from the *i*’th neuron will contribute **w**_*i*_ to the network output, while a spike from its ‘antineuron’ counterpart will have a contribution of −**w**_*i*_ to network output, but **w**_*i*_ itself may be positive or negative.

Formally, we define the population framework in terms of the update equations:

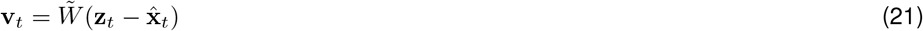

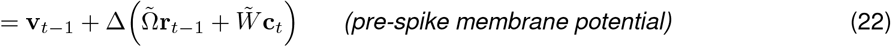

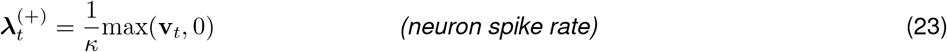

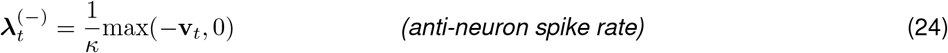

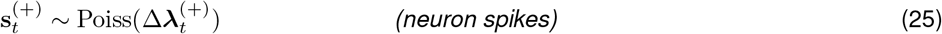

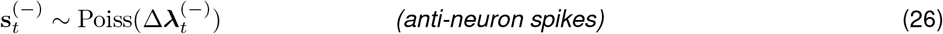

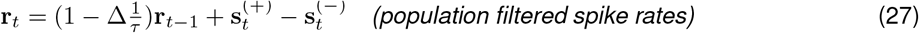

where 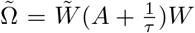 are the coupling weights from **r**_*t*−1_ to the pre-spike membrane potentials. This differs from the standard BSN framework in that spikes, rather than being driven by deterministic threshold crossing, arise from a Poisson process with conditional intensity 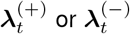. If **v**_[*i*]*t*_ is negative, then 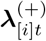 is set to zero, and the corresponding anti-neuron’s firing rate is positive. The voltage updates and spiking resets are identical to the local and BSN models.

Figure 6 shows a comparison of local and population-level frameworks. The population model achieved higher accuracy than the local model for both the one-dimensional and two-dimensional dynamical systems, although at the expense of an increased number of spikes. However, the performance of the local model depends on the parameters of the firing rate nonlinearity, and we could have increased accuracy by increasing steepness *α* and maximal firing rate *F_max_*.

**Figure 6:**
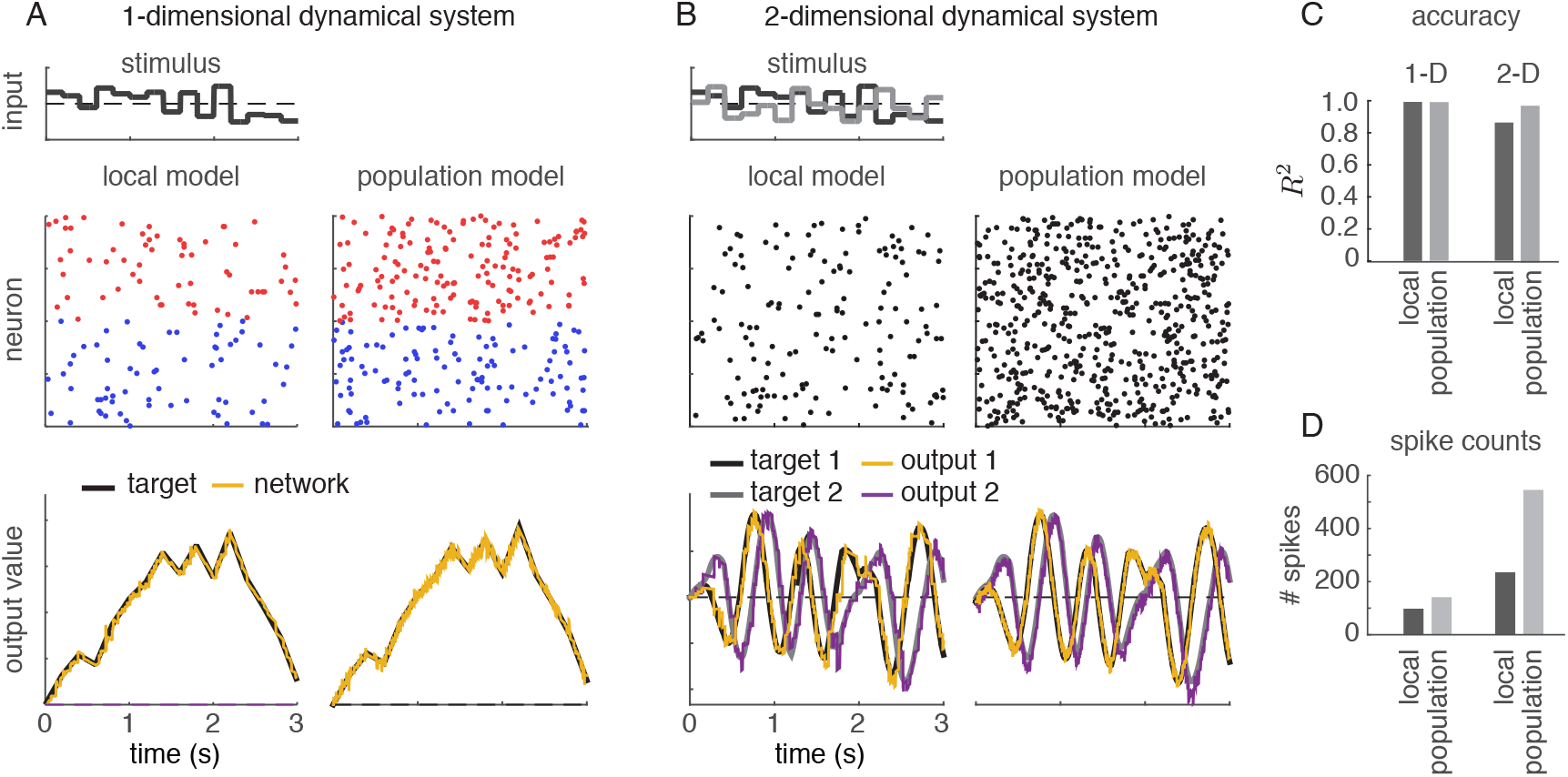
Simulations of the local and population frameworks implementing a 1D and 2D dynamical system. (**A**) The target was a 1dimensional integrator: *ẋ*(*t*) = *c*(*t*). Left side shows spikes and outputs from local Poisson model, while right side shows spikes and outputs for population Poisson model. As in previous figures, red dots indicate spikes from neurons with positive output weights, blue dots indicate spikes from neurons with negative weights. (**B**) The target was a 2-dimensional oscillator *ẋ*_1_(*t*) = −*ẋ*_1_(*t*) − 10*x*_2_(*t*) + *c*(*t*); *x*_2_(*t*) = 10*x*_1_(*t*) − *x*_2_(*t*) + *c*(*t*). For the population model, the time window for computing expected spike count was *κ* = 5ms (50 time bins). Weights were randomized to be positive or negative in either dimension, such that neurons are no longer divided into strictly positive- or negative-weight groups. **(C)** Accuracy (*R*^2^) of the two models for 1D and 2D systems. (**D**) Number of spikes emitted by each model during simulations.

## 5 Incorporating synaptic time delays

The original BSN model relies on near-instantaneous synaptic communication between neurons due to the fact that all neurons in the population reset immediately after a spike in any neuron. A more realistic model would require that synaptic inputs arrive only after a brief synaptic delay; only the reset of a neuron’s own membrane potential following a spike could be considered instantaneous.

To test the robustness of the two Poisson BSN frameworks introduced above, we altered synaptic currents to incorporate a synaptic delays between neurons. In the revised model, spike trains and filtered spike trains received by other neurons are updated only after a synaptic delay *d*. Thus, if neuron fires at time *t*, it resets its own membrane potential in the next time bin, but we will update spike trains and filtered spike trains received by other neurons only at time *t* + *d*.

To compensate for synaptic delays, we altered the network dynamics so that membrane potential reflects the network error extrapolated *d* time steps into the future:

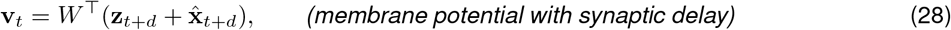

where **z**_*t*+*d*_ = exp(−*Ad*)**z**_*t*_ represents the network target at time *t* + *d*, computed by solving the homogeneous differential equation for **z**_*t*+*d*_ given an initial condition of **z**_*t*_ and linear dynamics matrix *A*, and 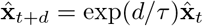 represents the network output at time *t* + *d* given by assuming passive decay of the membrane potential. Note here that exp(−*Ad*) denotes matrix exponential, and that this is equivalent to using exponential Euler integration to compute future values of **z** and 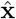 (see Methods for details).

Under these revised dynamics, a membrane potential at time *t* represents the extrapolated error between true and desired network output at time *t* + *d* instead of the instantaneous error at time *t*. This is equivalent to minimizing an objective function (eqs. 4 or 19) defined in terms of **x**_*t*+*d*_ and 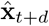 instead of **x**_*t*_ and 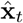. For the population-level Poisson framework, the encoding weights 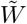 replace *W*^⊤^ in (eq. 28).

Figure 7 shows an analysis of the accuracy of the local and population-level Poisson frameworks with synaptic delays. For both models, a 1ms synaptic delay does not have pronounced effects on the spiking activity or the quality of the read-out (Fig. 7A-B). Fig. 7D shows the *R*^2^ values as a function of time delay for both frameworks. We compare it against a theoretical upper bound on the coding accuracy (in black), which is a consequence of the exponential Euler approximation. To determine this bound, we integrated the target dynamics with and without exponential Euler integration and calculated R^2^ for all values of *d*.

**Figure 7:**
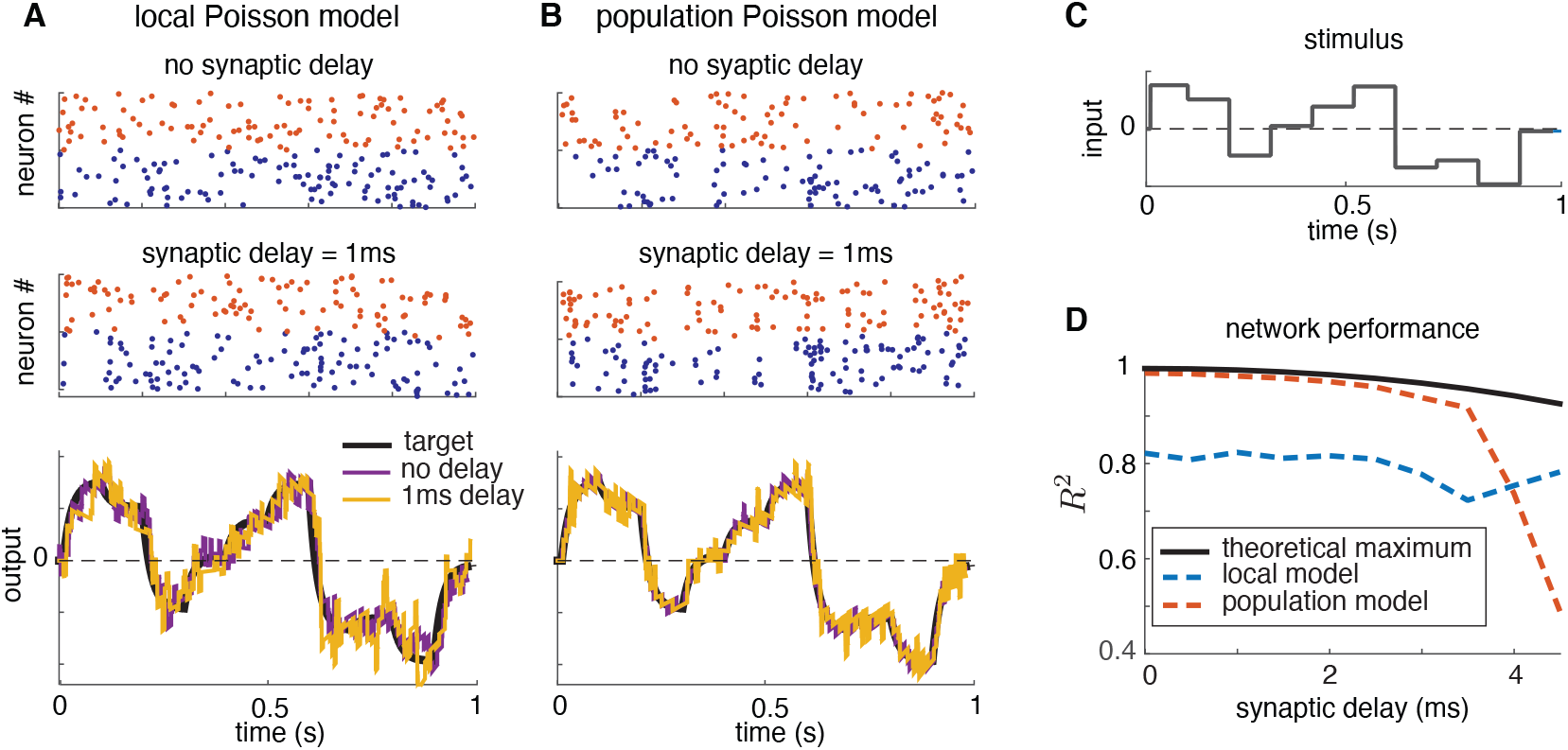
Illustration of local and population conditionally Poisson BSN frameworks with synaptic delays. **(A)** Spike trains simulated from the local Poisson framework implementing a 1D perfect integrator, both without (top) and with a 1-ms synaptic delay (middle). The network output accurately tracked the target variable for both models (bottom). As before, red/blue spike trains indicate neurons with positive/output weights. **(B)** Analogous plots for population Poisson framework. **(C)** Stimulus used for simulations shown in A and B. **(D)** Coefficient of determination (R^2^) computed using 50 simulations of each framework. Black trace indicates the maximum possible R^2^ value that could be obtained given the exponential Euler integration rule for computing the future target at time *t* + *d*.

For the local framework, the *R*^2^ value is lower than in figure 6 because the parameters were chosen to make the network more robust to synaptic delays. Otherwise, requiring high precision with synaptic delays results in ping-ponging, similar to the higher error regime shown in figure 5E. By contrast, the population framework can maintain high levels of accuracy for a large range of synaptic delays.

## 6 Cross-correlations

Lastly, we compared the statistics of spike trains generated by the original BSN with those of the local and population Poisson BSN models. Figure 8 shows the cross-correlations of spike trains generated by a network of forty neurons implementing a one-dimensional integrator, *ẋ*(*t*) = *c*(*t*), with a white-noise stimulus *c*(*t*). Local and population Poisson BSN models both enforced a synaptic delay of 1 ms.

**Figure 8:**
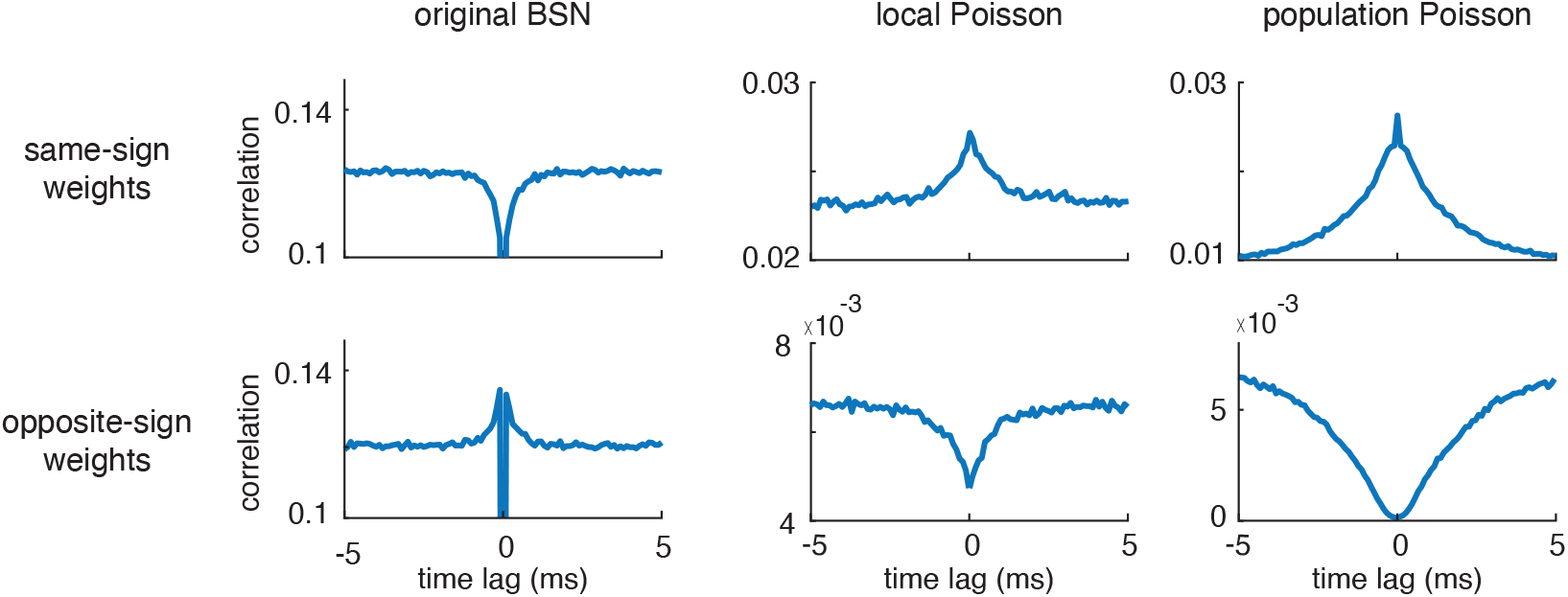
Cross-correlations for the original BSN, local Poisson and population Poisson BSN models with synaptic delay. Top row shows average correlations across pairs of neurons with the same sign output weight (i.e., both positive or both negative). Bottom row shows average correlations across pairs of neurons with opposite-sign output weights (i.e., one positive and one negative neuron). The original BSN network exhibits negative correlations between neurons with the same sign, and positive correlations between neurons with opposite sign. The local and Population Poisson models show the opposite pattern, which more closely resembles correlations found in neural populations in (e.g.) visual cortex.

To compute cross-correlations, we divided the neurons into positive-output and negative-output groups. We then computed average within-group (positive-positive and negative-negative) and across-group (positive-negative) cross-correlations. These curves show substantial differences between the original BSN model and the two Poisson models. First,cross-correlations of the original BSN model are 0 at lag zero,due to the rule that only one neuron can spike in a single time bin. More importantly,the within-group correlations for the BSN model exhibit a trough at time zero, meaning that neurons with the same output weight are anti-correlated. Conversely, across-group correlations exhibit an increase at small lags, meaning that neurons with opposite sign output weights are more likely to fire together in a small time window.

This relationship is at odds with correlations in both retina and visual cortex, where studies have reported that correlations are highest for neurons with similar tuning, and lowest for neurons with dissimilar tuning [34, 38–40]. By contrast, the local and population Poisson models successfully recapitulate this pattern of correlations, with a peak in the cross-correlations between pairs of neurons with the same sign weights, and a trough for pairs of opposite-sign neurons. Cross-correlations from these models also exhibit no trough at zero due to the lack of a rule prohibiting simultaneous spiking. Thus, cross-correlations represent an additional dimension of biological plausibility of the proposed Poisson frameworks.

## 7 Discussion

In this paper, we have highlighted a shortcoming of the balanced spiking network (BSN) paradigm, namely the requirement of near-instantaneous communication between neurons, which arises from the fact that a spike in any neuron causes an instantaneous reset of membrane potential in all other neurons. In practice, the BSN model is often implemented with the additional rule that only one neuron can spike in a single time bin. When synaptic delays are introduced, or multiple spikes are allowed per bin, the model easily enters a ping-ponging regime in which the network output overshoots and undershoots the target output on alternating time bins.

To address this problem, we proposed two extensions to the BSN model that incorporate conditionally Poisson spiking. Our proposed models both preserve the readout structure of the original BSN, in which a linear combination of exponentially filtered spike trains approximates a linear dynamical system of interest, and the desirable coding characteristics, such as E/I balance. However, they both replace of “hard threshold” integrate-and-fire spiking of the original BSN with a spiking process governed by an instantaneous spike rate or conditional intensity.

In the “local” Poisson BSN framework, the conditional intensity arises from passing the membrane potential through a sigmoidal nonlinearity. The accelerating phase of this nonlinearity is consistent with nonlinearities observed in neural data [41–44] and closely resembles the exponential nonlinearity commonly used in generalized linear modeling analyses [33, 34], while the saturating phase is consistent with saturation in real neural firing rates.

In the “population”Poisson BSN framework, the conditional intensity is obtained by setting the vector of expected spike counts to the least-squares solution for the total output error. This model differs from the original BSN in that the encoding weights, the linear mapping from output error to membrane potential, uses the pseudo-inverse of the decoding weights, 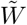, whereas the original BSN used the transpose *W*^⊤^. This change ensures that the spike rate of each neuron takes account of how many other neurons in the population have similar decoding weights, so that the expected spike count across the entire population in some finite time window compensates optimally for the output error. These modifications make both frameworks robust to both parameter settings and synaptic delays on realistic time scales (1-3ms).

### 7.1 Related work

Our paper is not the first to address the issue of instability in the BSN. Recent work from Koren and Denève [45] examined the use of penalties on spiking to reduce ping-ponging (referred to in that paper as “up states”). We found that this strategy required fine-tuning and succeeded in a relatively narrow parameter regime compared to the solutions we proposed here. Other work has argued that oscillations in the brain activity may arise from BSNs with synaptic delays, suggesting that a substantially damped form of ping-ponging may be a signature of efficient computation in neural circuits [46].

Recent literature has explored a variety of other extensions and applications of the BSN framework, including nonlinear dynamical systems and the learning of synaptic weights [47, 48], synaptic plasticity rules [49], and biological extensions like finite timescale synapses [50] and synaptic delays [45]. The BSN framework has also been adapted to other computational problems such as probabilistic computation [51] and sensory adaptation [52].

The topic of balanced networks has also received considerable attention outside the specific BSN framework introduced by Boerlin et al. [1]. Balanced networks have been proposed as a substrate for working memory [53, 54], probabilistic inference [55, 56], and the control of complex movements [57]. Excitatory-inhibitory balance is also a key topic in the mathematical theory of neural circuit dynamics, where it has been proposed as an explanation for the correlations found in large-scale population activity [58–60]. Finally, a rich literature has focused on the training of spiking neural networks in more general supervised and reinforcement learning settings, where the objective involves task performance or can only be evaluated at the end of a trial [61–65].

Our work also connects to a rich literature on point process models of neural spike trains. The local Poisson framework draws direct inspiration from the work of Plesser and Gerstner [32], which sought to approximate a noisy integrate-and-fire model with an inhomogeneous Poisson process via the so-called “escape-rate approximation”, which refers to the instantaneous probability of noisy membrane potential crossing threshold in a small time window. Subsequent work on the spike response model [36, 66–69] and Poisson generalized linear model [33–35, 70–72] further explored the connection between integrate-and-fire and conditionally Poisson spike train models. The latter are sometimes referred to as “soft-threshold” integrate-and-fire model [73], making the local Poisson model a natural extension of the original BSN model.

### 7.2 Future challenges

Although our proposed frameworks are a step in the direction of biological plausibility, there remain a variety of open challenges. One such challenge is the development of neurally plausible learning rules and weight patterns. The network we proposed has a static weight matrix with all-to-all connectivity. A more realistic model would allow for sparse connectivity, sign constraints forcing neurons to be purely excitatory or inhibitory, and plausible learning rules that allow weights to change over time as a function of reward signals. The mathematical treatment of learning in the context of the BSN has proven difficult, although there have been successes learning the fast, slow and feed-forward weights through non-local, supervised, control theoretic approaches [47, 48]. The probabilistic formulation of the Poisson BSN frameworks makes implementing local, Hebbian plasticity rules more tractable, as it opens up the possibility of applying unsupervised learning techniques from traditional machine learning methodology. Finally, we hope to implement time delays in a more principled way, as in [50], and to address the biological plausibility of ‘anti-neurons’ in the population framework.

A second challenge is the incorporation of nonlinear dynamics. Although the original BSN model was designed to implement linear dynamical systems, it is well known that a wide variety of neural computations are nonlinear. Recent work has proposed an extension of the BSN framework to nonlinear dynamics [31, 47]; combining this approach with conditionally Poisson spiking therefore represents a promising avenue for future work.

Finally, the conditionally Poisson extensions we have proposed provide new opportunities for applying the BSN framework to the interpretation and analysis of real neural data sets. Both the original BSN model and ours assume access to the precise spiking patterns of all the neurons in a population, but real neural recordings typically record only a small fraction of the neurons in a population. Previous work has shown that latent BSN dynamics can be recovered from spike trains in the fully observed case [51]. Other work has discussed the recovery of Poisson generalized linear models from partial recordings [74, 75]. This motivates the development of new methods for identifying balanced network dynamics and computations from partially observed data sets, which may offer fundamental insights into spike-based computation in the brain.

### 8 Methods

#### Exponential Euler integration

Exponential Euler integration is a method for solving first-order differential equations of the form

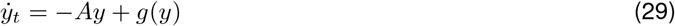

where −*Ay* is a linear term and the nonlinear terms are grouped in *g*(*y*). For equations without nonlinear terms, it above can be solved exactly from time 0 to a later time *t* as

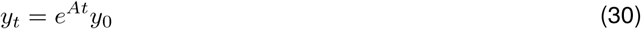

We use this method to approximate the value of a differential equation at a later time, *t′*, by approximating the value of a function *y_t_* at time *t* + *t′* as

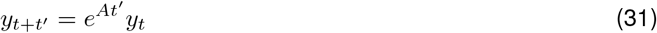

where the accuracy of the approximation decreases as *t′* increases.

#### Simulation parameters

For all simulations, time is measured in seconds and *dt* = 0.1ms. All other simulation parameters are shown in the table below.

Figure 2 was generated using the parameter settings described in Boerlin *et al*, fig. 1C (included below for comparison). The cost terms are *μ* = 10^-6^ and *ν* = 10^-5^ and the voltage decay constant is τ_v_= 20. The noise added to the voltage dynamics and the stimulus was Gaussian with *σ_v_* = 10^-3^ and *σ_c_* = 0.01,respectively. When enforcing the constraint that one neuron should spike per time bin, we selected the neuron with the highest voltage above threshold. Another option with similar results would be to select randomly between the ones above threshold.

**Table 1:** Simulation parameters

| Figure | N | $W$ | $\alpha$ | $F_{max}$ | $F_{min}$ | $\kappa$ | $A$ | $\tau$ | $d$ |
| --- | --- | --- | --- | --- | --- | --- | --- | --- | --- |
| 2 | 400 | $\pm 0.1$ | N/A | N/Ax | N/A | N/A | 0 | 10 | 0 |
| 4D, BSN | 1 | 1 | N/A | N/A | N/A | N/A | 0 | 20 | 0 |
| 4D, GLM | 1 | 1 | Varies | 50 | 0 | N/A | 0 | 20 | 0 |
| 5 | 400 | $\pm .1$ | 1000 (ACD) 100 (B) | 100 (ABD) 10 (C) | 1 (A-C) 0 (D) | N/A | 0 | 10 | 0 |
| 6A-B | 400 | $\pm .2 + \text{noise } (\mu = .4)$ | 80 | 1 | 0 | 50 | 0 | 10 | 0 (A) 1ms (B) |
| 6C-D | 400 | $\pm .2 + \text{noise } (\mu = .4)$ | 80 | 1 | 0 | 50 | $\begin{pmatrix} -1 & -10 \\ 10 & -1 \end{pmatrix}$ | 10 | 0 (C) 1ms (D) |
| 7 | 200 | $\pm 0.02 + \text{noise } (\mu = .01)$ | 500 | 100 | 0 | 25 | -50 | 5 | 0 and 1ms |
| 8 | 40 | $\pm 0.0025 + \text{noise } (\mu = .01)$ | 500 | 10 | 0 | 20 | 0 | 2 | .1ms |

As a reminder, *N* is the number of neurons in the network and *W* is the vector of read-out weights. For the local framework, *α* is the slope of the exponential nonlinearity, *F_max_* is the saturation, and *F_min_* is the minimum firing rate. For the population framework, *κ* is the time window over which the network minimizes the error. Finally, *A* is the dynamics matrix of the linear dynamical system, *τ* is the decay time constant of the filtered spike trains **r**, and *d* is the time delay.

A software implementation of Poisson BSNs, along with code to re-generate all simulation figures shown in the manuscript, is available at https://github.com/pillowlab/PoissonBalancedNets.

#### Cost Terms

The original BSN model objective function incorporated two additional cost terms to penalize spiking: a quadratic cost term *μ* and the linear cost term *ν*. These terms encouraged the network to use fewer spikes and to distribute spiking more evenly across neurons with large and small output weights. We did not incorporate these in our derivation for clarity, but they are included in our simulations of the BSN.

Including both cost terms into the derivation in Section 2, equation (4) becomes

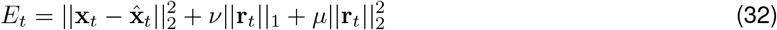

and the voltage and threshold equations become

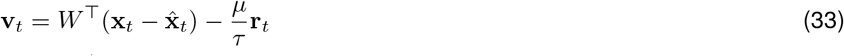

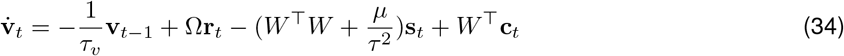

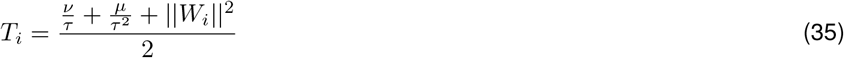

The linear cost term (*ν*) is proportional to the L1 norm of **r**(*t*), or 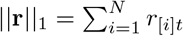. This cost term penalizes the network’s total activity. The quadratic cost term (*μ*) limits individual neuron firing rates, forcing a spread of activity across all neurons in a population. Over time, the network transfers activity from precise, costly neurons with high firing rates to imprecise, larger weighted neurons to maintain a compromise between efficiency and accuracy of the read-out. Boerlin *et al* also include a voltage leak term for biological realism.

In our Poisson models, we did not observe the ping-pong effects described in Boerlin *et al* for the range of parameters we considered, so we don’t need cost terms for network stability. For the local Poisson framework, the cost terms can be included when *α* and *F_max_* are high enough to cause ping-ponging.

